# Using Google Earth to improve management of threatened limestone karst ecosystems in Peninsular Malaysia

**DOI:** 10.1101/048397

**Authors:** Thor-Seng Liew, Liz Price, Gopalasamy Reuben Clements

## Abstract

Biodiversity conservation is now about prioritisation, especially in a world with limited resources and so many habitats and species in need of protection. However, we cannot prioritise effectively if historical and current information on a particular habitat or species remains scattered. Several good platforms have been created to help users to find, use and create biodiversity information. However, good platforms for sharing habitat information for threatened ecosystems are still lacking. Limestone hills are an example of threatened ecosystems that harbour unique biodiversity, but are facing intensifying anthropogenic disturbances. As limestone is a vital resource for the construction industry, it is not possible to completely halt forest degradation and quarrying in developing countries such as Malaysia, where 445 limestone hills have been recorded in the peninsula to date. As such, there is an urgent need to identify which hills must be prioritised for conservation. To make decisions based on sound science, collating spatial and biological information on limestone hills into a publicly accessible database is critical. Here, we compile Malaysia’s first limestone hill GIS map for 445 limestone hills in the peninsula based on information from geological reports and scientific literature. To assist in conservation prioritisation efforts, we quantify characteristics of limestone hills in terms of size, degree of isolation, and spatial distribution patterns and also assessed the degree of habitat disturbance of each limestone hill in terms of buffer area forest degradation and quarrying activity. All this information is stored in a KMZ file and can be accessed through the Google Earth interface. This product should not be viewed as a final output containing basic limestone hill information. Rather, this database is a foundational platform for users to collect, store, update and manipulate spatial and biological data from limestone hills to better inform decisions regarding their management.

## Introduction

In a world with limited resources, biodiversity conservation is becoming more about prioritisation – making difficult choices on which habitats of an ecosystem or which populations of a species to conserve (Myers et al., 2000; Brooks et al., 2006). However, conservation priorities cannot be identified unless ecosystems are quantified and mapped (Olson et al., 2001; Naidoo et al., 2008). With advancements in remote sensing technology, different ecosystems, such as forests, rivers and lakes, can now be detected and be mapped effectively. Ecosystems that have yet to be adequately mapped include limestone hills in karstic areas, which cover about 11% of the Earth’s surface (Williams, 2008).

Limestone hills in the tropics are regarded as “arks” of biodiversity as they contain high levels of species endemism (Schilthuizen, 2004; Clements et al., 2006; Chua et al., 2009; Liew et al., 2014). Quarrying of limestone hills represents the most direct threat to its biodiversity as it causes irreversible destruction of habitats (Clements et al., 2006; Schilthuizen & Clements, 2008). In addition, degradation of forest on and around limestone hills can also result in negative impacts on its biodiversity (Schilthuizen et al., 2005). In fact, many species that are endemic to limestone hills are already extinct or at the brink of extinction (Liew et al., 2014; Liew, 2015). As limestone hills will continue to be exploited by the cement industry, there is an urgent need to prioritize limestone hills for conservation (Vermeulen and Whitten, 1999; Clements et al., 2006).

In Malaysia, karstic areas with limestone hills are vital components of the country’s geoheritage (Hussain et al., 2008). Many of these hills were connected and belong to the same geological formation or bedrock whereas some of these hills are lenticular and never connected to any other hills (Paton, 1961). These hills distribute unevenly across Peninsular Malaysia, and almost all the hills (> 95%) are found in the four states of Kelantan, Pahang, Perak, and Perlis. However, information on the location of limestone hills in the country remains is scattered in the literature.

The most reliable information sources have been geological reports that include maps illustrating the location of limestone hills in a given area (see Supplementary Information 1). Several rather comprehensive gazetteers are also available, but in most cases, they provide only names, approximate coordinates, and descriptions of the limestone hills without complete maps of where they are located (e.g. Price, 2001; 2015). To date, there are no publicily accessible, free, complete, reusable, easily editable and scientifically rigorous maps of limestone hills available for scientific research and management.

Remote sensing technologies have had moderate successes in detecting limestone hills in tropical regions where the limestone hills are subject to more intensive continuous weathering (de Carvalho et al., 2013; Theilen-Willige et al., 2014; Kobal et al., 2015). The efficiency of the detection from remote sensing approaches depends on the scale, sensor and the geomorphology of the limestone (Siart et al., 2009; Rajendran & Nasir, 2014). However, many of the limestone hills in tropical regions such as Malaysia are very small, often without visible geomorphological features, and are covered by forest – this poses a challenge to distinguish them from regular hilly forests using remote sensing alone.

As it is not possible to protect every limestone hill in Malaysia due to economic demands, a conservation prioritisation exercise will have to be conducted to improve the management of this threatened ecosystem. Before this can be done, we first need baseline scientific data for each limestone hill in terms of its physical and threat characteristics. Subsequently, people can add information on their geological, biodiversity, archaeological and geological importance. Unfortunately, such data is only available for a small number of hills and even so, the different datasets have yet to be integrated into a single database. One of the main challenges to form such a database is the fact that there is a lack of resources (i.e. time and money) and expertise to gather such scientific data. Another fundamental problem is that there is no general reference scheme to identify limestone hills; different studies can use different names for the same hill, which could then hamper data integration.

In view of these issues, we develop a working limestone hills GIS map for Peninsular Malaysia. We first compile localities of known limestone hills in this region from scattered publications and digitised them as a KML (Keyhole Markup Language) format file. With this database established, we can then conduct a preliminary conservation prioritization exercise for limestone hills by quantifying their: 1) physical parameters based on physical parameters such as size, degree of isolation, and spatial distribution patterns; and 2) threat parameters based on the presence/absence of quarrying and degree of habitat disturbance in terms of buffer area forest degradation. Finally, these outputs will be saved as a KMZ file for public access via the Google Earth interface.

## Material and methods

### Limestone hills mapping

We manually compiled limestone hill data from scattered literature and systematically transformed them into an accessible GIS database using a multi-level approach similar to GIS remote sensing approaches (e.g. Theilen-Willige et al., 2014). First, we extracted localities of limestone hills from 61 publications, which were mainly geological references that included good quality geological maps. We did not include hills that are only presented by coordinates. We excluded the limestone hills on the offshore islands for this research.

Second, we marked these hills in Google Earth as polygon placemarks and annotated them with their reference source in the “description” field of the placemark. Due to its, rich temporal repository of high resolution images, most of the hills were visible on Google Earth. Ironically, recent forest loss and monoculture crop plantations around limestone hills have also made them more prominent and readily identifiable. However, several limestone hills were very small and covered by forest – this can make them indistinguishable from the surrounding forest. As such, we used the shape of the hills from relevant maps to verify their location.

Third, we added additional hills that are omitted from the literature based on unpublished field notes. Lastly, digitisation of the limestone hills was peformed directly in the Google Earth by tracing the outline of the limestone outcrop as a polygon. At the same time, the code/name of the hill used by those references were tabulated in the description field of the polygon properties, for example: [Source: Gobbett, 1967]<Hill name: Bukit Biwah>. For limestone hills that were already listed in the most comprehensive gazeteer for limestone hills in Peninsular Malaysia (Price 2015), we used its limestone hill reference number and hill name for the name of the polygon. For remaining hills not listed or cannot be verified in Price (2015), we used prefix of “mykarst” and follow by unique numbering (i.e. 001 onwards). The digitised outline and reference information of each hill were saved in a single KML file to be analysed and modified later.

### Quantifying limestone hill physical parameters

Figure 1 illustrates the steps involved in this process. First, we converted the KML file consisting of 445 limestone polygons into a polygon limestone hill shapefile (*.shp), which has a Universal Transverse Mercator coordinate system. Next, we quantified the limestone size, degree of isolation and distribution patterns from this shapefile by using QGIS version 2.8.3 – Wien (QGIS Development Team, 2015). Last, we plotted the limestone size against the degree of isolation to explore the irreplacibility limestone hills – larger and more isolated hills would be considered more irreplacable based on the central tenets of island biogeography. All the GIS data can be found in Supplemental Information 2.

**Figure 1.**
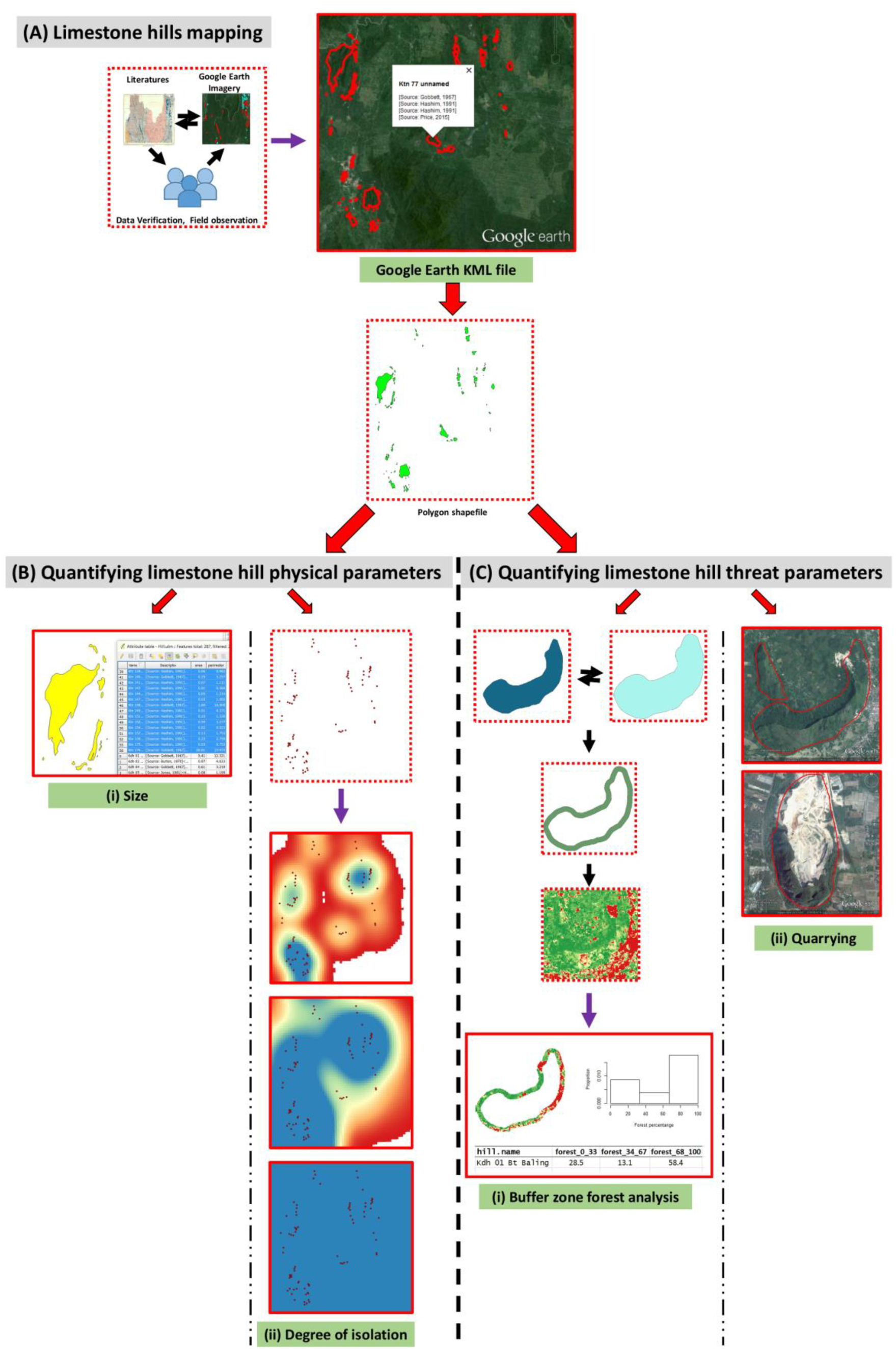
Analytical flowchart for this study. (A) limestone hills mapping; (B) quantifying limestone hill physical parameters – (i) size and (ii) degree of isolation; (C) quantifying limestone hill threat parameters “ (i) buffer zone forest status and (ii) quarrying.

To calculate the size of each limestone hill (i.e. area; km^2^), we used the “Geometry$area” functions of the “Field calculator” in the attribute table of the shapefile. A series of histograms with different bin sizes were produced to examine the distribution of the limestone hill sizes in Peninsular Malaysia.

To calculate the degree of isolation of each limestone hill, we converted the hill polygon shapefile to point shapefile so that each hill was represent by its centroid point by using “polygon centroid” function. After that, we generated limestone density raster layer with a cell size of 50 meters by using “Heatmap analysis” for 10km, 25 km and 50 km radius respectively. Next, we used “Add grid values to points”, in SAGA version 2.1.4 (Conrad et al., 2015), to extract the density values for the 10 km, 25 km and 50 km radius’ density raster layer for each of the limestone hill (centroid) to the limestone hill point shapefile’s attribute table.

The three density values of each limestone hill were used as degree of isolation of each limestone hill in three different spatial contexts. For example, a limestone hill with raster value of 4 in the 10 km heatmap raster layer means that there are three other hills within 10 km radius of the limestone hill. The small spatial context (e.g. 10 km search radius) is useful for visually identifying clusters for isolated hills in Peninsular Malaysia, whereas the larger spatial contexts are useful to produce a generalised density map where clusters could not be easily identified. By doing this, an extremely isolated hill should have lower density values for all spatial contexts, while an isolated limestone cluster should have higher value in the small spatial context but lower value in larger spatial contexts. We then explored limestone hills’ density values for all spatial contexts by using histogram to identify isolated single hill and limestone clusters, in which several separate hills can be situated in close proximity to each other, but are far apart from other limestone clusters. All the plots were made using R version 3.2.2 (R Core Team. 2015).

### Quantifying limestone hill threat parameters

To determine the quarrying disturbance of limestone hills, each hill of the KML polygon file were inspected visually from Google Earth for the presence/absence of quarrying activity. Due to resource constraints, we do not ground-truth to verify whether the quarrying sign of a limestone hill indicates an abandoned or active quarry.

To assess the the forest status at the buffer zone of limestone hills, we used the buffer zone polygon shapefile of each limestone hill and published forest status maps from literatures. For forest status maps, we used the Landsat datasets of Hansen et al. (2013), which provided tree canopy cover in the year 2000. The layer has grid size of 1 arc-second (^~^30 meters) and each grid cell has percentage value in the range 0–100 for all vegetation taller than 5 m height. In addition, we also used forest gain layer and forest loss layer of Hansen et al. (2013; accessed Dec 2015). The details of these layers are available at http://earthenginepartners.appspot.com/science-2013-global-forest/download_v1.2.html. We converted the coordinate system of these layer to UTM (Timbalai 1948 / UTM zone 50N - EPSG:29850 - EPSG.io). We aware that there are other forest maps that derived from different remote sensing imageries with different resolutions, such as Sophie et al. (2010), Miettinen et al. (2012), and Shimada et al. (2014). However, large forest extent maps created using simple algorithms are not without weaknesses and discrepancies (Dong et al., 2012; Tropek et al., 2014). In fact, using a high resolution forest map of Peninsular Malaysia is critical for a buffer zone analysis involving relatively small features such as limestone hills. Until this kind of map available, we felt it was necessary to conduct an exploratory analysis of the forest condition in the limestone buffer zone using the most suitable map for our purpose (i.e. Hansen et al., 2013).

Next, we generated the 250 m buffer zone of each hill based on polygon limestone hills shapefile. After that, the area size of this buffer zone was calculated. All the three forest status layers were cropped by this buffer zone. For the forest cover layer, cell values ranging from 1-100 were summarised into three bins in histogram, namely, (1) 0-33; (2) 34-67; and (3) 68-100, for each hill. On the other hand, the forest gain layer and forest loss layer were recalculated to produce a new layer that represent both net forest loss and gain in the buffer zone of each hill. Lastly, we explored the forest status of each hill by plotting the proportion of ‘good’ forest (value 68-100) against the proportion of net forest loss in the buffer zone of each hill. All these analyses were done in R environment by using Packages “raster” (Hijmans, 2015), “rgdal” (Bivand et al., 2015)” and “rgeos” (Bivand & Rundel, 2015).

### Create limestone GIS database in the form of KMZ file

In order to create a publicly accessible and user-friendly limestone GIS database, we incorporated the digitised map, references and limestone physical and threat parameters into a single KMZ file that can be accessed via Google Earth interface. We used the same the KML that already with digitised limestone hills outlines and reference information as a template. After that we summarised each limestone hill’s physical and threat parameter in one overall graph that consists of six plots and maps (Figure 2). Lastly, we updated the script of KML template to include the graph by a short custom written R script (Supplementary File 2).

**Figure 2.**
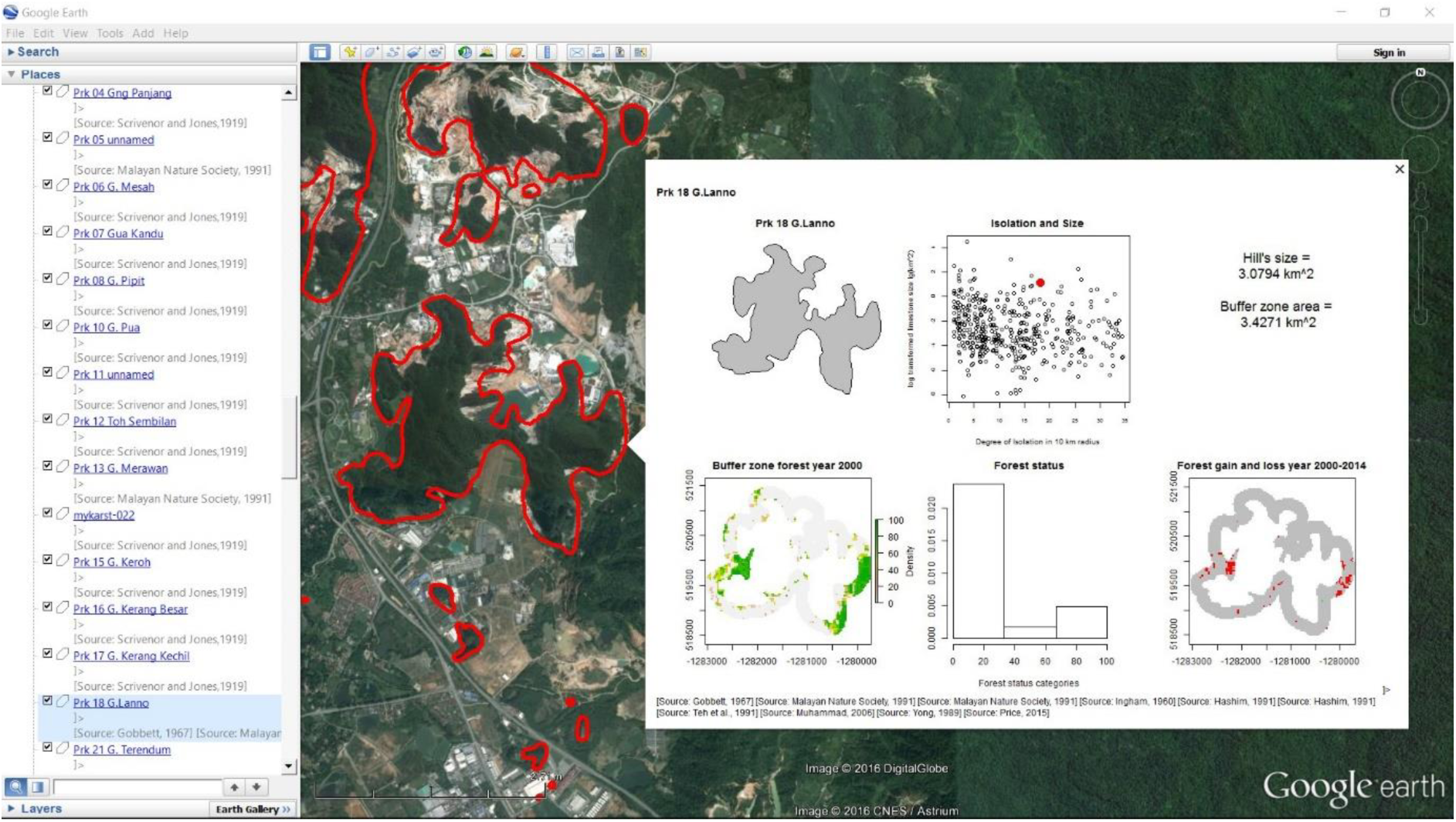
A screenshot of Peninsular Malaysia limestone hills GIS database open in Google Earth. The outline of each limestone hill is shown, and when selected (“click”), a pop out window shows the name, an overall graph and the references. The overall graph consists of: Top row (left to right) – a figure of limestone hills outline, a scatterplot shows the size and isolation of the hill in the context of other hills in Peninsular Malaysia, and hill and buffer zone area size; Bottom row (left to right) – map of forest status (i.e. Hansen et al., 2013) in buffer zone in year 2000, a histogram that summarised forest status in buffer zone in year2000, and a map shows the forest net gain and net loss in buffer zone between year 2000 – 2014.

## Results

### Limestone hills mapping and GIS database

In total we digitised 445 hills via Google Earth based on the literature (Figure 3). Of Peninsular Malaysia’s total land surface of 130598 km^2^, only 280 km^2^ (^~^0.2%) is covered by limestone hills. The frequency of each limestone hill mentioned in the references were shown in Supplementary Information 1: Figure S1).

**Figure 3.**
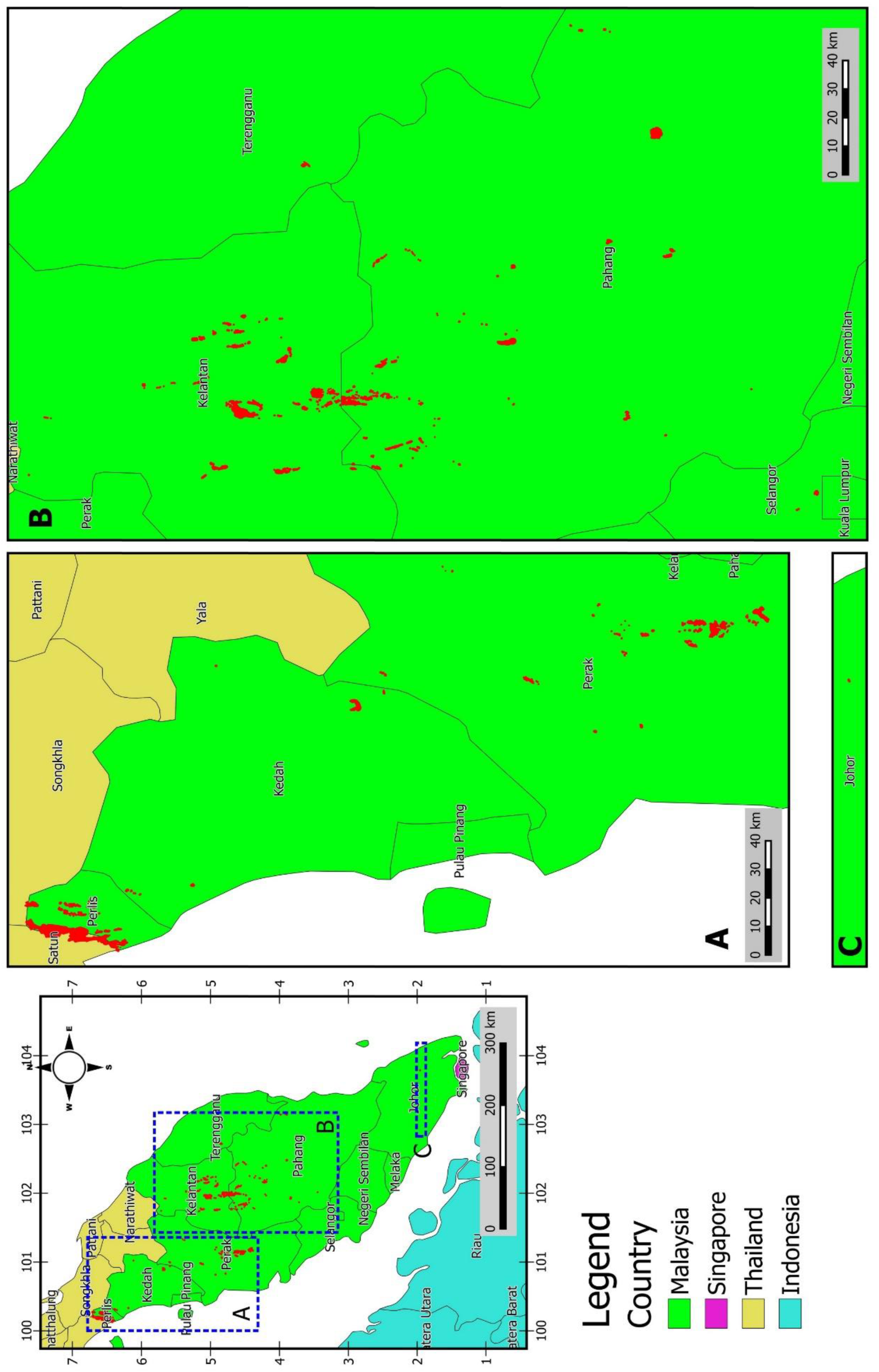
Maps of limestone hills in Peninsular Malaysia.

During the collation of the spatial data, we found two interesting patterns. First, 81 of 445 limestone hills were recorded only once from the references that we surveyed; for the majority of hills, the karstic area of a particular hill was actually covered by more than one references. Second, many of the limestone hills that were recorded numerous times revealed inconsistencies among hill names, including well-studied limestone hills (e.g. “Prk 27 G. Datuk”, “Prk 29 Gunung Lang”). The original KML polygon file of these digitised hills can be found Supplementary File 3.

The state of Kelantan has the most number of hills – 149 (33.5%), followed by Pahang – 124 hills (27.9%), Perak – 93 hills (20.9%), and Perlis – 60 hills (13.5%). The remaning 19 hills (4.25%) are distributed at Kedah – 12, Selangor – 3, Terengganu – 3, and Johor – 1 (Figure 3). A GIS database in a single KMZ with a small file size (34.6 MB) was created. It contains basic information on the physical and threat parameters of each of the 445 hills that can be accessed via Google Earth. In addition, advanced users can create a newer version of this database (KMZ file) by adding or deleting hill outlines in the original KML file according to the needs of the user (e.g. some people may just want to conduct a conservation prioritization exercise for hills in a single state). Our analyses can be repeated for different user scenarios by following the analytical workflow described above and R script (Supplementary File 2).

### Limestone hill physical parameters

Histograms in Figure 4 show the distribution of limestone hill sizes for three different series of interval. The histogram is strongly skewed to the left. Around 90% of the limestone hills in Peninsular Malaysia are smaller than 1 km^2^. There are only three hills larger than 10 km^2^, namely, Prs 64 Wang Ulu – 85.30 km^2^, Ktn 176 Batu Baloh – 21.2 km^2^, and Phg 77 Bukit Mengapur – 11.8 km^2^. Another 17 hills have size ranging from 2 km^2^ – 10 km^2^, and 22 hills have size ranging from 1 km^2^ – 2 km^2^. Around 60 % of the hills are smaller than 0.1 km^2^ and a further 14 % of the hills smaller than 0.01 km^2^.

**Figure 4.**
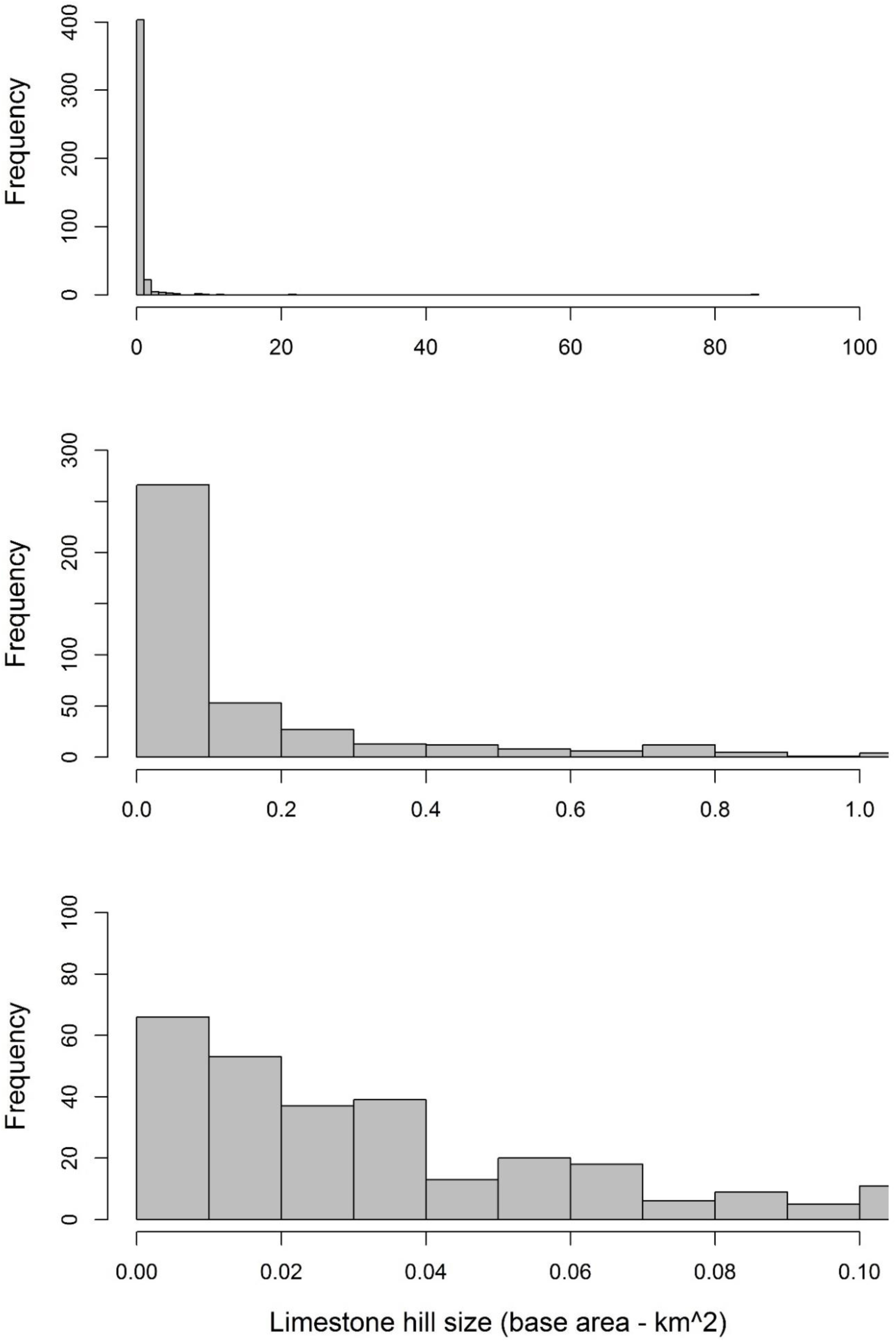
Histograms show the distribution of the size of limestone hills in Peninsular Malaysia. (A) Histogram of all 445 hills with bin’s size of 1 km^2^. (B) Histogram of those hills smaller than 1 km^2^ with bin’s size of 0.1 km^2^. (C) Histogram of those hills smaller than 0.1 km^2^ with bin’s size of 0.01 km^2^.

Figure 5 shows the distribution of the degree of isolation for each hill in three different spatial contexts. More than half of the limestone hills is near to seven other limestone hills within 10 km (62%). The most isolated hills for each spatial context were shown in Table 1. There are two extremely isolated hills each of the three contexts with only single hills within 10 km, 25 km, and 50 km radius, namely, "mykarst-034" and "mykarst-057". In addition, density maps of 10 km search radius indicate that there are 22 limestone clusters that are relatively more isolated from the other hills (i.e. at least 10 km away from other nearby cluster), and 15 of these clusters do not consist more than 5 hills within each cluster (Figure 6).

**Figure 5.**
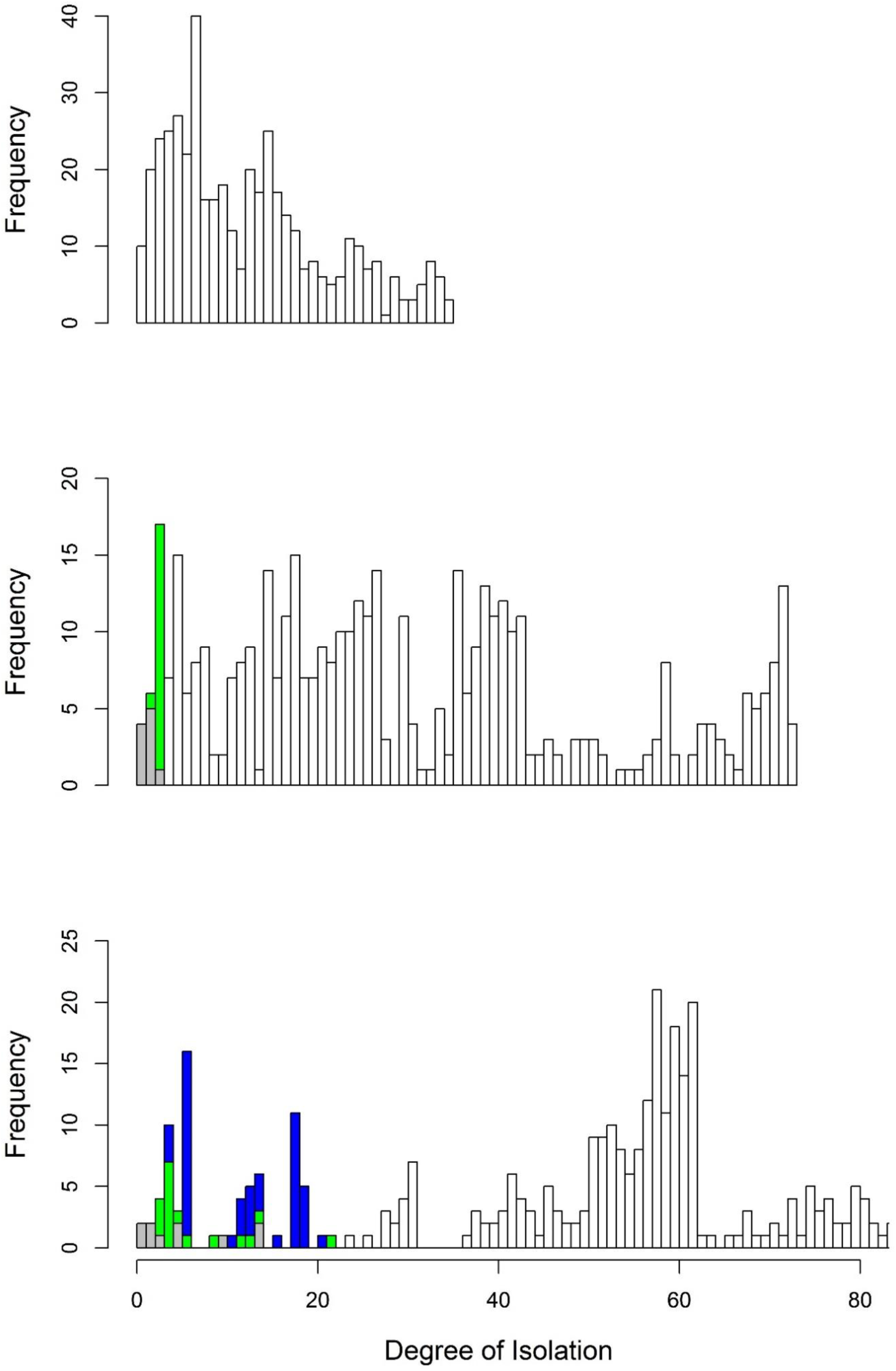
Histograms show the degree of isolation (i.e. density values) in three different geographical contexts: (A) Number of adjacent hills within 10 km radius of each hill. (B) Number of adjacent hills within 25 km radius of each hill. (C) Number of adjacent hills within 50 km radius of each hill.

**Figure 6.**
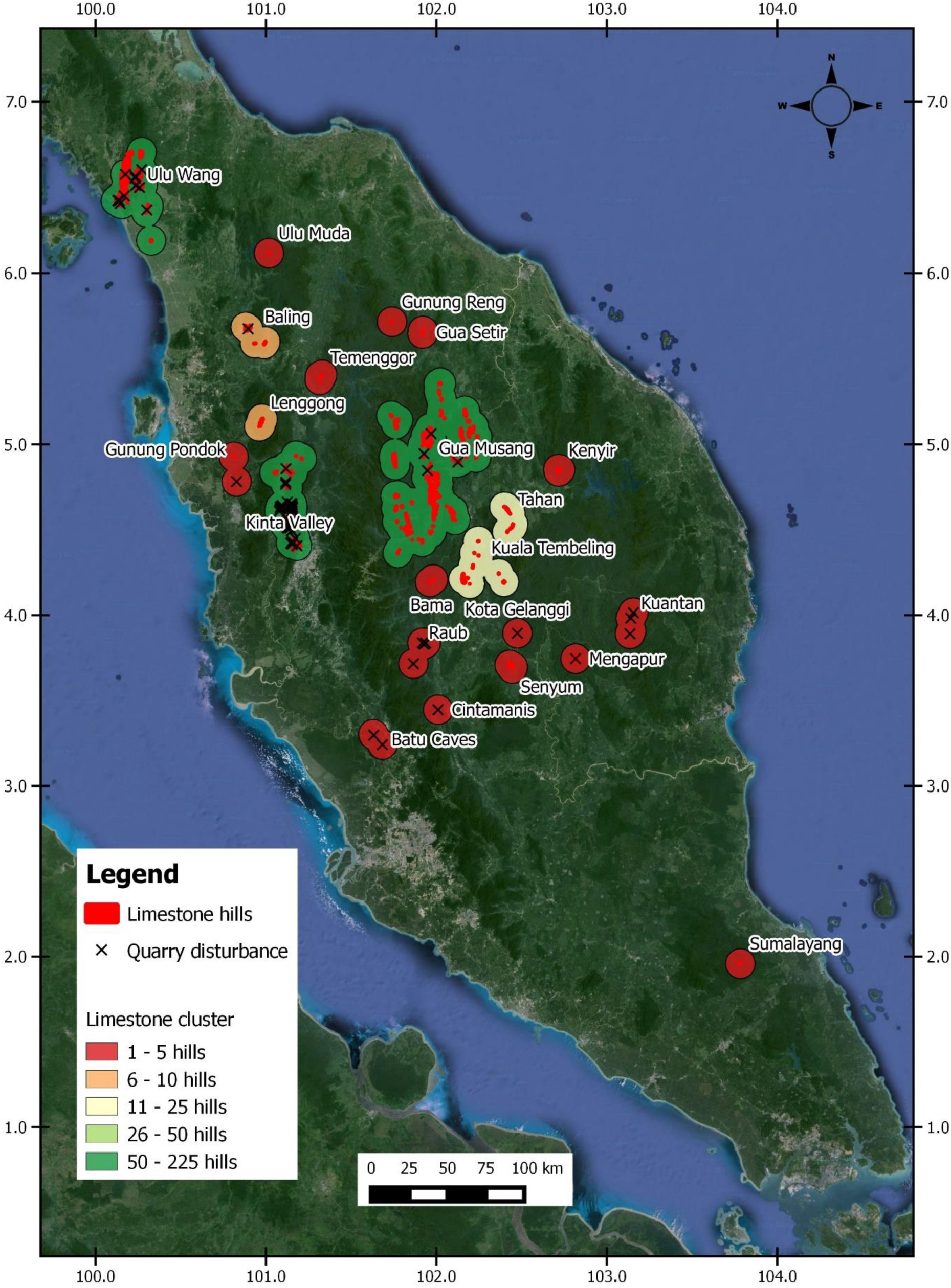
Map of limestone hills clusters in Peninsular Malaysia shows that number of limestone hills and the quarrying activities in each cluster. Imagery ©2016 Data SIO, NOAA, U.S. Mavy, NGA, GEBCO, Landsat, Map data ©2016 Google

### Limestone hills threat parameters

A total of 73 of 445 limestone hills (16%) in Peninsular Malaysia have signs of operational or historical quarrying activities (Figures 6 and 7). There was at least one hill completely quarried away – “Phg 72 Bt Panching”. It appears there is no clear association of whether or not the hill was quarried and limestone physical parameters (Figure 7).

**Figure 7.**
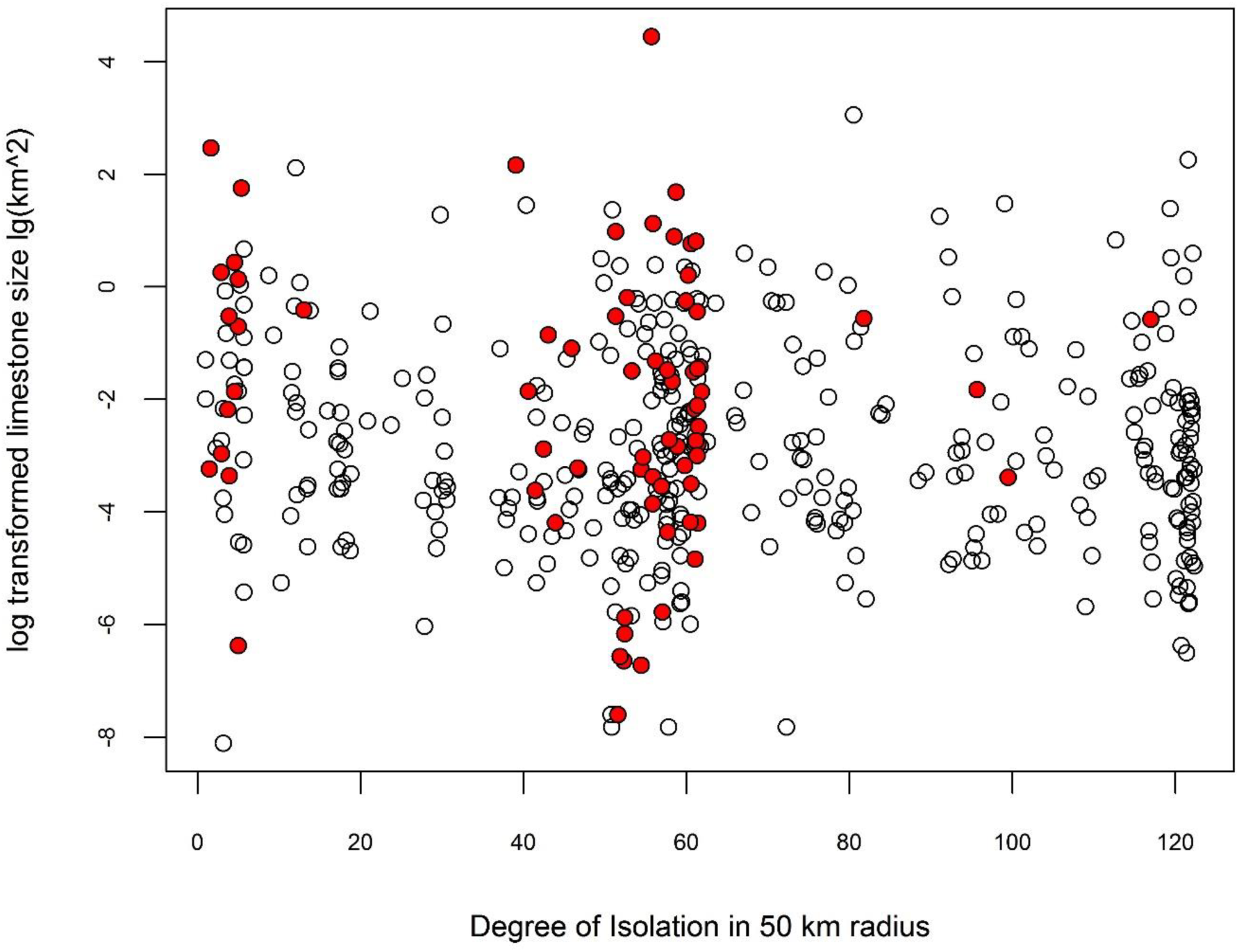
Size and degree of isolation of all the limestone hills in Peninsular Malaysia. Red closed circles are the hills with quarrying disturbance.

In terms of buffer zone status, half of the limestone hills appear to have good forest coverage (i.e. more than 80 % of the buffer zone still with the forest coverage value above 67; Figure 8). About 10% of the limestone hills have buffer zone forests that were highly degraded (i.e. each with more than 80 % buffer zone having forest coverage value below 34).

**Figure 8.**
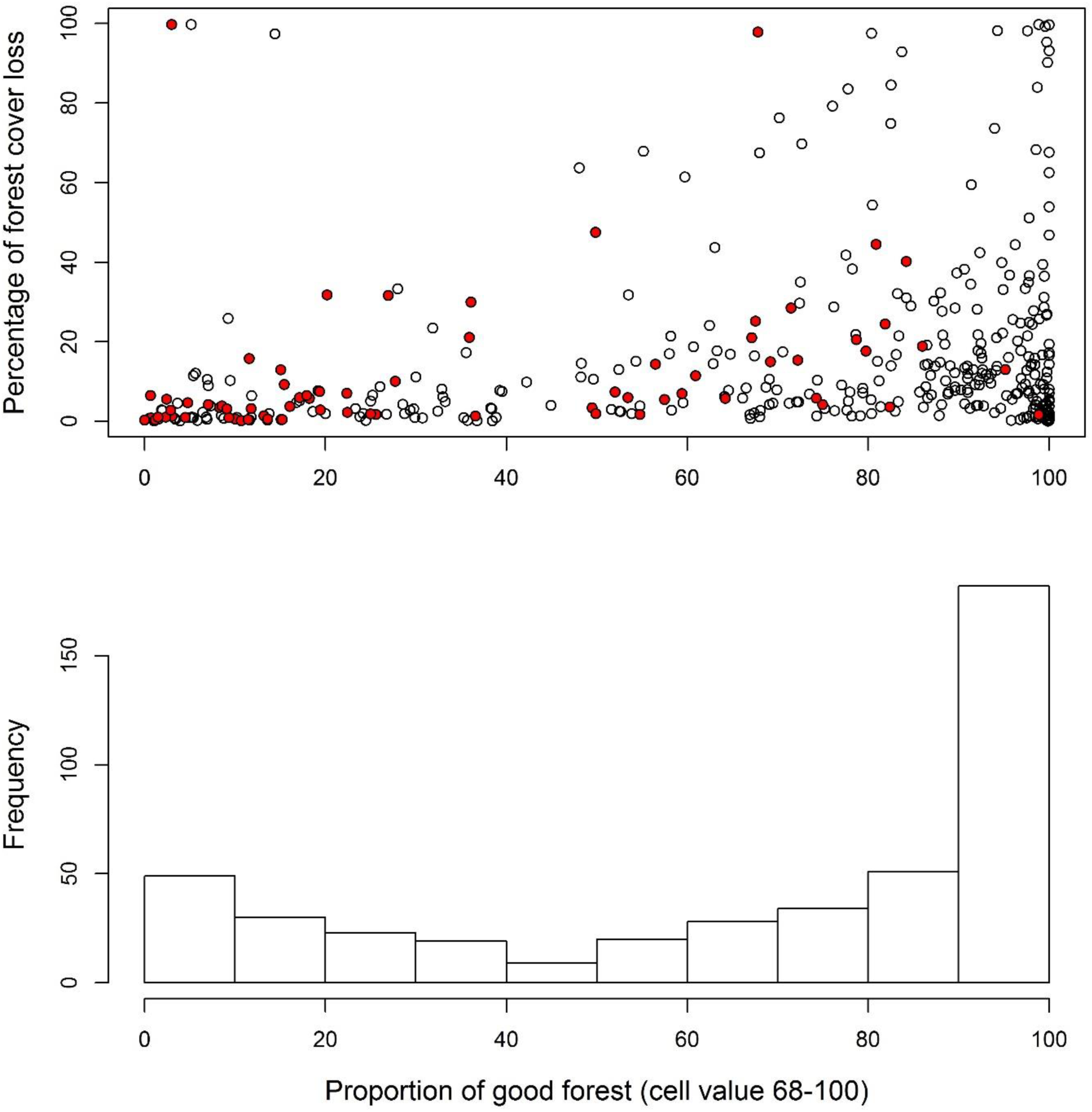
Forest status in buffer zone. (A) Proportion of forest with cover value (68-100) in year 2000 and percentage of forest loss from year 2000 – 2014 in the buffer zone of each limestone hill. Red closed circles are the hills with quarrying disturbance. (B) Histogram of the proportion of forest cover value (68-100) remains in the buffer zone of all the limestone in Peninsular Malaysia.

## Discussion

Our GIS database, which consists of 445 limestone hills marked on Google Earth with their names and reference sources, represents the first up-to-date, comprehensive and accessible source for limestone hills in Peninsula Malaysia and possibly the tropics.

Many of the limestone hills were recorded several times independently, while some hills were recorded only once. Our records, however, require further ground verification, especially hills that were recorded only once. As already noted by Davison and Kiew (1990) and Price (2001), the exact number of limestone hills is difficult to determine due to the ambiguous definition of a single hill. We define a single hill as one that is completely separated from nearby hills aboveground. There were previously recorded hills that we did not include in our database due to uncertainty; for example, we recorded 149 hills in the state of Kelantan, whereas, Davison and Kiew (1990) estimated 210 and verified 120 of them; and Price (2001) recorded 234 hills. Nevertheless, we believe our GIS database can be the first reference point for future verification. Furthermore, the analytical workflow that we used in this study can be repeated to generate up-to-date database when new hills are found or new forest cover layers are available in the future.

We suggest that our reference code assigned to each limestone hill be used by future users when they report their findings from limestone hills. Previously, biodiversity sampling at limestone hills often reported the name of limestone hill along with GPS coordinates. However, there have been incidents of inconsistent use of limestone hill names, which can create confusion especially when the reference source of the hill name is not provided. In addition, GPS coordinates taken by different users with different datum can result in discrepancies of tens or hundreds of meters, which can be an issue when there are many hills close to one another. Most importantly, the inconsistent use of hill names and coordinates can hinder the sharing and integration of information from various scientific publications. Hence, we hope this database can be a reference platform for information collected from limestone hills throughout Peninsular Malaysia.

### Limestone hill physical parameters

Limestone hills in the tropics are considered de facto islands as they are surrounded by non-calcareous substrata (Clements et al., 2006). Hence, the biodiversity richness and endemism patterns on these hills are mainly shaped by the same ecological and evolutionary processes as on islands. Two main island characteristics, size and isolation, have been shown to be the main determinants of endemism and geographically structured genetic patterns for species on limestone hills (Vermeulen & Whitten, 1999; Clements et al., 2008; Latinne et al., 2012; Haskell & Pan, 2013; Gao et al., 2015).

In terms of limestone hill sizes, our database indicates that the majority of hills in Peninsular Malaysia are smaller than 0.1 km^2^, of which a quarter are smaller than a football field (0.01 km^2^). Currently, very little is known about the biological importance of these small hills and the minimum size required to support significant levels of biodiversity – this is a very important information gap to be filled before important management decisions can be made regarding their ‘irreplacebility’. When such information is available, for example, a threshold line can be plotted in Figure 7 to determine conservation priorities.

In terms of limestone hill isolation, this characteristic can be assessed from the aspect of a single hill, or a cluster that consists of a number of adjacent hills. Such information is very important as it is not easy to formulate a management plan for all 445 hills in Peninsular Malaysia at this moment. One way to address this issue is to group these unevenly scattered hills into a number of manageable working clusters using a systematically quantitative and qualitative approaches. In this study, we identified the clusters based on the degree of isolation when a group of hills is less than ten kilometres away from one another, and at the same time, the cluster is approximately ten kilometres away from other limestone clusters. Nevertheless, the limestone hill clusters identified by this approach need to be refined with an integrated and systematic approach using quantitative and qualitative information on their geological formation, extent of cave networks, biodiversity and biogeography.

When this is done, a more coordinated and rigorous research and conservation program can be designed within a limestone cluster or between clusters. A limestone cluster could consist of a number of lenticular hills that are not on a single bedrock or number of hills that are on the same limestone bedrock. Hence, a number of limestone hills can be grounded in a limestone cluster, just like a number of islands are grouped as an archipelago. Even high-level conservation decisions such as the process of nomination and inscription of limestone hills for world heritage has been done ad hoc, which has led to a suboptimal representation of the limestone hill values without considering the ‘irreplacebility’ of these hills in the overall context of hills in a region (Williams, 2008).

### Limestone hill threat parameters

Only a few limestone clusters in Malaysia have been accredited World Heritage status, namely, Mulu (Sarawak), Niah (Sarawak), and Lenggong (Perak). In addition, there are a few more clusters that have been protected as a National Park such as Langkawi Islands, and those in Kuala Tahan, Kuala Tembeling and Kenyir. However, the majority of limestone hills in Malaysia do not have protected area status and are vulnerable to anthropogenic disturbances.

Generally, limestone hills have sparser vegetation covers, especially for tiny hills that are overlaid by thin soil layers. This type of habitat is extremely sensitive to anthropogenic disturbances such as deforestation and quarrying (Vermeulen & Whitten, 1999; Tuyet, 2001; Clements et al., 2006; Zhang et al., 2011). In addition to the size and isolation factors mentioned above, the degradation of the forest and quarrying of limestone hills could change the biodiversity pattern and genetic structures of the organisms on limestone hills (Schilthuizen et al., 2005; Flavenot et al., 2015).

One immediate management intervention that can be adopted to better protect limestone biodiversity is to preserve a wide forested buffer zone around the base of limestone hills (Vermeulen & Whitten, 1999; Davison & Kiew, 1990; Schilthuizen et al., 2005). Davison and Kiew (1990) suggested a forested buffer zone of at least 200 m around the hill which to prevent burning, to provide shaded habitat and to protect stream system. Our analysis of buffer zone forest status seems to suggest that most of the forest in buffer zones are still in reasonably good conditions in year 2000, with relatively low forest cover loss in the past decade. However, as mature oil palm plantations also have high forest cover values in Hansen et al. (2013), we caution that the forest cover in the buffer zones of many limestone hills in our analysis could be actually oil palm plantations, and this analysis needs to be repeated when an improved forest cover data is available in the future.

We found that many of the limestone hills with very poor forest conditions in buffer zones have been quarried. As compared to deforestation, quarrying is more destructive and its impact is irreversible (Clements et al., 2006). The most common practice in the industry is to completely quarry away a limestone hills and then move to the adjacent hills. It seems that there has not been a dramatic increase in the number of quarries operating – Price (2001) reported that there was a total of 62 limestones quarries in 1975. To date, we have detected 73 limestone quarries. However, the main cause for concern is that there are ongoing quarrying activities are occuring in seven of the highly isolated and small (< 5 hills) clusters, namely, Raub, Gunung Pondok, Mengapur, and Kuantan (Figure 6). Hence, the companies mining these extremely ‘vulnerable’ hills should be engaged with urgently in order to develop biodiversity and conservation plans.

## Conclusion

Our current version of database should be seen as a first reference point to provide quantitative and qualitative information of limestone hills in Peninsular Malaysia that can assist users to verify and build upon (e.g. Foody, 2014). We acknolwedge that our database may not have had a rigorous ground-truthing of data quality to be considered a ‘final’ map because there are many hills that still require further verification. Nevertheless, limestone hills in different states within Peninsular Malaysia can now be prioritised for conservation in the context of systematic conservation planning (Margules & Pressey, 2000) by using our physical and threat parameters as proxies of their ‘irreplacebility’ and ‘vulnerability’, respectively. We hope that this user-friendly karst database can better facilitate the communication and sharing of spatial information, especially among ordinary citizens to increase their concern for their natural environment (Butler, 2006). Furthermore, our database could encourage stakeholders of the limestone hills, including scientists, industries and policy-makers, to enrich the database with more information on their archaeological, speleological, geological, biodiversity, cultural, and economic importance.

## Acknowledgement

We thank Lahiru S. Wijedasa for the discussion about the problems in mapping limestone hills in Peninsular Malaysia, Annebelle Kok for assisting to scanning the geological map, Menno Schilthuizen for financial sponsor to purchase some of the literatures. We also thank ## reviewers and editor for their constructive comments on the earlier draft of this paper.

## Author Contributions

TSL conceived and designed the experiments, performed the experiments, analysed the data, wrote the paper, prepared figures and/or tables, reviewed drafts of the paper.

PL performed the experiments

RGC conceived and designed the experiments, wrote the paper, reviewed drafts of the paper.

## Table

**Table 1.** Top isolated 40 limestone hills in Peninsular Malaysia.

## Supplemental Information

https://dx.doi.org/10.6084/m9.figshare.3153181.v1

**Supplemental Information 1.** References used in digitalisation of limestone hills.

**Supplemental Information 2.** Limestone hills GIS file (KML format).

**Supplemental Information 3.** GIS data and R script for data analysis.

